# Bidirectional Generative Adversarial Representation Learning for Natural Stimulus Synthesis

**DOI:** 10.1101/2023.10.17.562789

**Authors:** Johnny Reilly, John D. Goodwin, Sihao Lu, Andriy S. Kozlov

## Abstract

Thousands of species use vocal signals to communicate with one another. Vocalisations carry rich information, yet characterising and analysing these complex, high-dimensional signals is difficult and prone to human bias. Moreover, animal vocalisations are ethologically relevant stimuli whose representation by auditory neurons is an important subject of research in sensory neuroscience. A method that can efficiently generate naturalistic vocalisation waveforms would offer an unlimited supply of stimuli with which to probe neuronal computations. While unsupervised learning methods allow for the projection of vocalisations into low-dimensional latent spaces learned from the waveforms themselves, and generative modelling allows for the synthesis of novel vocalisations for use in downstream tasks, there is currently no method that would combine these tasks to produce naturalistic vocalisation waveforms for stimulus playback. In this paper, we demonstrate BiWaveGAN: a bidirectional Generative Adversarial Network (GAN) capable of learning a latent representation of ultrasonic vocalisations (USVs) from mice. We show that BiWaveGAN can be used to generate, and interpolate between, realistic vocalisation waveforms. We then use these synthesised stimuli along with natural USVs to probe the sensory input space of mouse auditory cortical neurons. We show that stimuli generated from our method evoke neuronal responses as effectively as real vocalisations, and produce receptive fields with the same predictive power. BiWaveGAN is not restricted to mouse USVs but can be used to synthesise naturalistic vocalisations of any animal species and interpolate between vocalisations of the same or different species, which could be useful for probing categorical boundaries in representations of ethologically relevant auditory signals.

## Introduction

Artificial intelligence is increasingly used to predict and better understand how neurons respond to natural stimuli. For example, evolutionary algorithms and convolutional neural networks (CNNs) can be used to identify stimuli that maximally drive single neurons in the auditory and visual cortices [***Chambers et al., 2014, Walker et al., 2019***]. Similarly, unsupervised neural networks employing learning rules grounded in biological principles are able to extract meaningful features from animal vocalisations which can provide context for the features discovered from neural recordings [***Ngiam et al., 2011, Kozlov and Gentner, 2016, Lu et al., 2023***]. Furthermore, CNNs have been shown to be effective in modelling auditory coding of complex, natural sounds, even generalising well to unseen neuronal data [***Arneodo et al., 2021***]. Such studies underscore the versatility and potential of deep learning models.

Natural stimuli such as animal vocalisations serve as a powerful tool to understand how the brain processes ethologically relevant sensory information, offering a more realistic perspective than simple artificial stimuli commonly used in experimental contexts [***David et al., 2004, Woolley et al., 2005, Kozlov and Gentner, 2016, Rose et al., 2021***]. It is well documented that mice and other rodents use a rich system of vocalisations at frequencies above the human hearing range (greater than 20 kHz). These vocalisations are referred to as *ultrasonic vocalisations (USVs)*. USVs are understood to be social signals that are produced in a variety of contexts, for example, adult male mice are known to produce USVs in the presence of female mice, and pups will emit them when separated from their mothers [***Gourbal et al., 2004, Haack et al., 1983***]. Neurons in the auditory cortex respond to these behaviourally salient sounds [***Liu et al., 2006, Liu and Schreiner, 2007, Carruthers et al., 2013, Lu et al., 2023***], which are typically emitted as sequences of discrete ‘syllables’, sometimes organised into song-like repeated phrases [***Holy and Guo, 2005***]. Much research has gone into analysing the structure of USV syllables and attempting to classify them into different types [***Coffey et al., 2019, Fonseca et al., 2021***]. However, there is no consensus on the number of syllable classes or how to classify them. Indeed, recent deep-learning based approaches to understanding the distribution of USVs seem to favour the idea that USV syllables cannot be clustered into discrete categories based on their discernible features but must be understood as varying continuously. Despite the efforts to analyse the structure of these vocalisations, there are few methods available to computationally and systematically generate these vocalisations [***Goffinet et al., 2021, Sainburg et al., 2019***].

By leveraging advances in deep learning, we developed a method capable of efficiently generating novel vocalisations. We show that these vocalisations are suitable naturalistic stimuli to probe the mouse auditory system: they drive neurons in the auditory cortex well and produce receptive-field features that predict neuronal responses to natural USVs just as well as features obtained using natural USVs do. Given that the mouse is a powerful animal model, with its plethora of genetic and behavioural tools, we believe that this method will be a valuable addition to the toolset for the study of sensory processing. In principle, given enough data for model training, our method can be used to produce, and interpolate between, vocalisations of any animal species, offering a rich source of stimuli for sensory neuroscience and other applications.

## Results

### Bidirectional WaveGAN

Our approach is based on WaveGAN [***Donahue et al., 2019***], a Generative Adversarial Network (GAN) which has been successfully used to generate realistic waveforms of human speech and birdsong. GANs consist of two neural networks: The generator, ***G***, which attempts to learn a mapping from a low dimensional latent space to the data domain, and the discriminator, ***D***, a binary classifier which attempts to distinguish between real and synthetic data generated by ***G***. The latent space provides a low dimensional encoding of the data that can be used for downstream tasks, or probed to explore to see what features the model has learned to encode. However, standard GANs do not come equipped with an ‘encoder’ that can map a data point to its latent vector representation, which limits its use as a representation learning tool. A class of architectures that do provide such capabilities are Variational Autoencoders (VAEs). VAEs consist of two neural networks, an encoder that maps data into a low-dimensional latent space, and a decoder that generates data from its learned latent representation. The benefit of using VAEs for representation learning is that, unlike GANs, they are bidirectional so not only can we generate data from points in the latent space but we can also map (real or synthetic data) into its latent representation. However, compared to GANs, VAEs are considered less capable of creating realistic synthetic data. For this reason, we opt to use the GAN modelling approach over a VAE architecture.

To address these shortcomings, we developed BiWaveGAN, a bidirectional GAN [***Donahue et al., 2016***] which includes an additional encoder network, ***E***, which learns to encode the data, mapping waveforms onto their latent representations. The output of the encoder network is incorporated into the loss function to ensure it learns an accurate encoding and is optimised jointly alongside the generator and discriminator. To our knowledge, this is the first reported work to apply bidirectional, encoding GANs to the problem of representing and synthesising realistic animal vocalisations.

Note that there are various other approaches to generative modelling that may perform well, such as autoregressive models (e.g., WaveNet [***Oord et al., 2016***] for audio generation) and diffusion models (state-of-the-art for image modelling) [***Dhariwal and Nichol, 2021***]—however, it is not obvious how to adapt such models for bidirectional representation learning.

In Figures 1 and S1 we present spectrograms of several samples drawn by the generator, alongside spectrograms of real USVs recorded from mice. The generator is able to produce a wide variety of vocalisations which indicates that no significant mode collapse has occurred. While the model’s output is varied, some syllables are generated better than others, with longer and more complicated syllables being poorly generated. This is likely because they are less common in the dataset so the model cannot learn to generate them as well as the more common shorter syllables. Despite being of high fidelity, the generated samples still possess distinguishable characteristics that differentiate them from real data. These characteristics manifest as artefacts present both audibly and visually in the spectrograms of the generated samples. In particular, when played back (slowed down by a factor of 16 to be audible) the samples do mostly achieve the distinctive whistling timbre of a USV, as well as recognisable shapes in spectrograms, but a repetitive stuttering background noise can be heard. These artefacts are likely the consequence of the generator’s upsampling procedure, and it remains unclear to what extent they would disappear upon further training.

**Figure 1.**
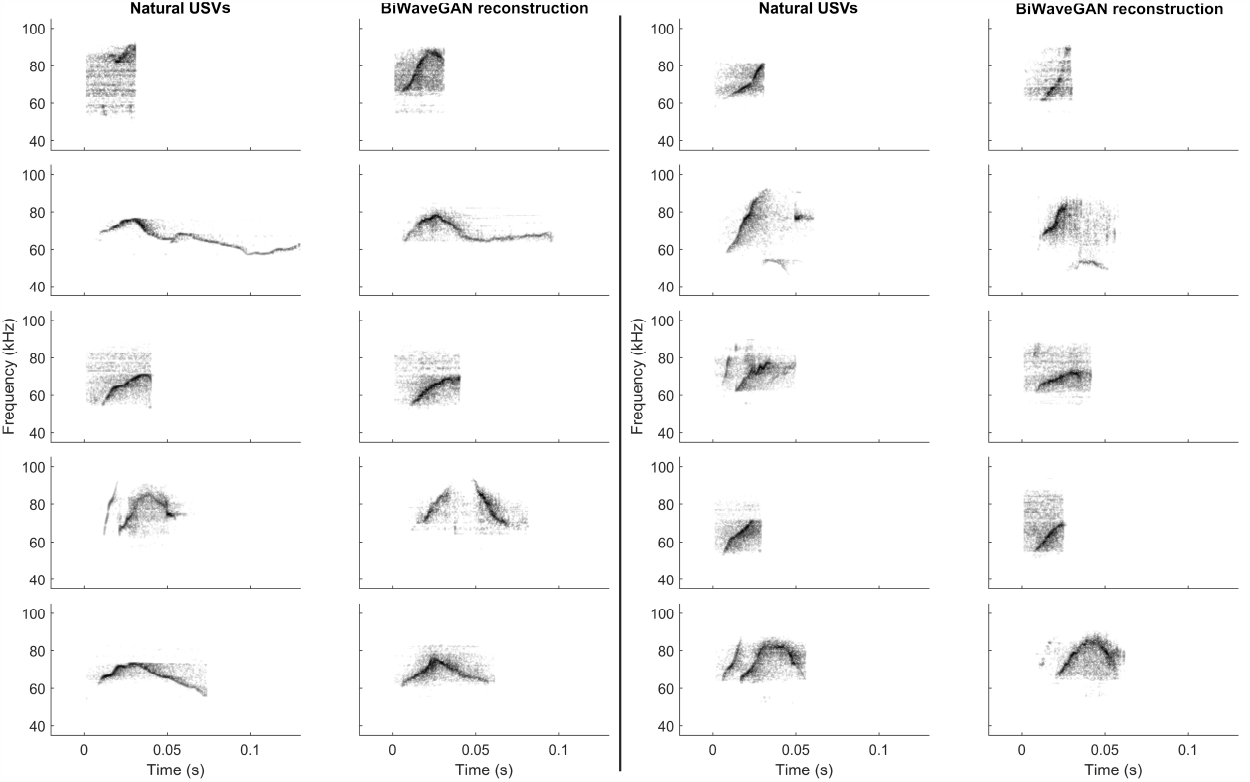
Example spectrograms of natural USVs and their BiWaveGAN-generated reconstructions. Spectrograms of natural USVs are shown in columns 1 and 3 with their corresponding reconstructions shown in columns 2 and 4. Reconstructions are produced by extracting the latent space representation of the corresponding natural USV with the encoder network, and then passing this vector into the decoder network to produce a synthetic USV.

Moreover, to evaluate the fidelity of the encoder and generator in their mutual inversion, we compare actual USVs from the dataset with their corresponding reconstructions, represented as ***G***(***E***(*x*)) where *x* denotes the test syllable. Spectrograms of these reconstructions are presented in Figure 1. The model is capable of reconstructing a wide array of vocalisations well, and even in instances where the real USV is not accurately reconstructed, the reconstruction looks realistic and displays similar temporal and spectral content when compared to the original.

Additionally, we explored the latent space further by examining 10-point linear interpolations between two points in the latent space. These interpolations allow us to assess the smoothness of the representation. The results of these interpolations are shown in Figure 2. While the reconstructions themselves are imperfect, it is clear that the model has learned to smoothly interpolate between USVs, even if they are very different from each other. Further, the intermediate waveforms look like reasonable USVs. This shows that the model has successfully learned a continuous latent representation of the data. The fact that the model is able to show us believable interpolations between USVs that would be considered categorically different shows that the syllables are not clustered. If the syllables were tightly clustered, we would not observe gradual change in interpolations but sharp changes as the representation ‘jumps’ from one class to the other.

**Figure 2.**
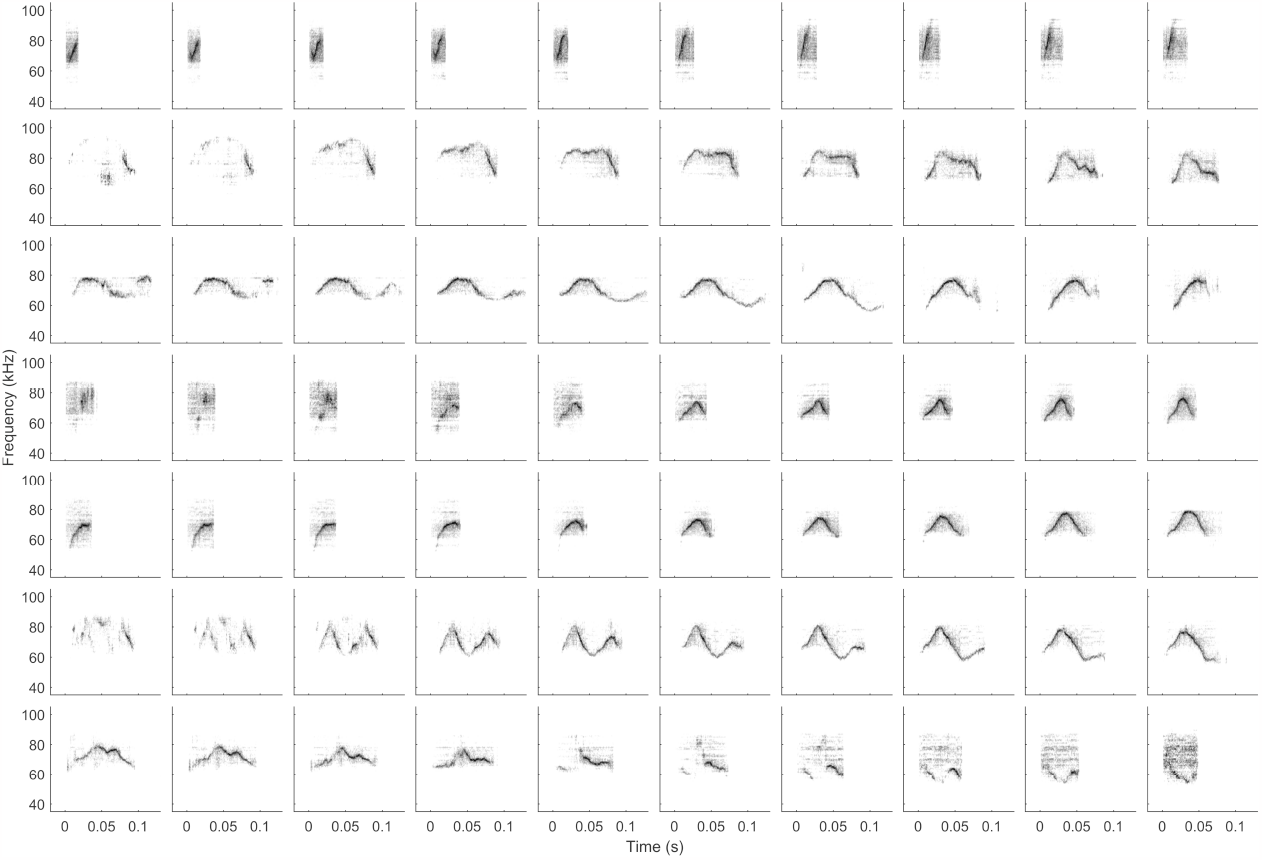
Example interpolations between pairs of USVs in latent space. Interpolations are produced by randomly sampling start and endpoints from the latent space, and then linearly interpolating eight intermediate values. These latent vectors are decoded to produce a sequence of ten synthetic USVs (one for each row in the image). As can be seen in the figure, linear interpolation in the latent space produces smooth variation in the USVs, even though the start and endpoint USVs look very different. This suggests that dissimilar USVs do not belong to discrete clusters.

### *In Vivo* Neural Responses to Reconstructed Stimuli

Spiking activity from a total of 46 single units was recorded from the right auditory cortex of 6 anaesthetised female C57BL/6J mice, during presentation of natural and BiWaveGAN-generated USVs. An example of one unit’s spiking activity in response to three syllables is shown in Figure 3. The power spectra of natural and reconstructed USVs were compared to ensure that any potential differences in neural response were not simply due to differences in the spectral profile of the stimuli (Fig. 4**A**).

**Figure 3.**
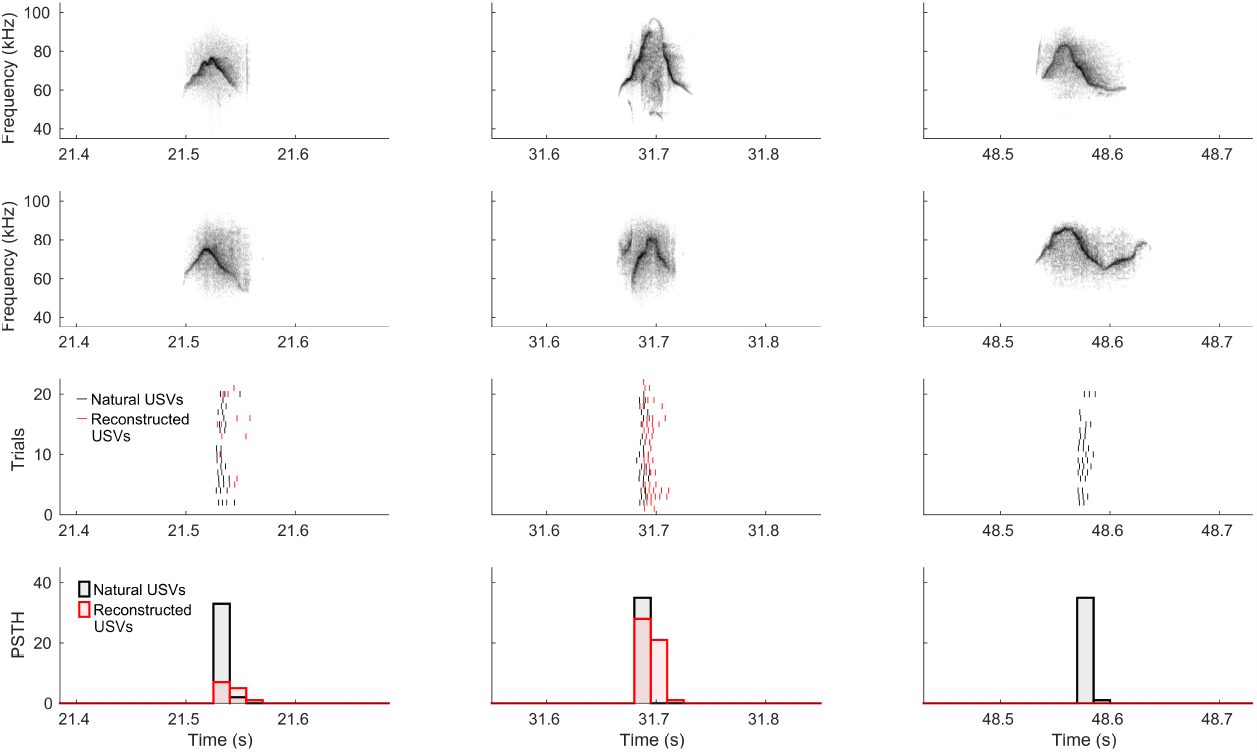
Example responses of a single unit to natural and BiWaveGAN-generated USVs. Spectrograms of natural and reconstructed USVs are shown in rows 1 and 2, respectively. Row 3 shows the spike responses of the unit during 22 repeated presentations of the natural USV stimulus shown in black, and the reconstructed stimulus shown in red. Corresponding peri-stimulus time histograms are shown in row 4. The first two pairs of syllables evoke similar responses within each pair, whereas the third syllable’s reconstruction evokes no response in this unit.

**Figure 4.**
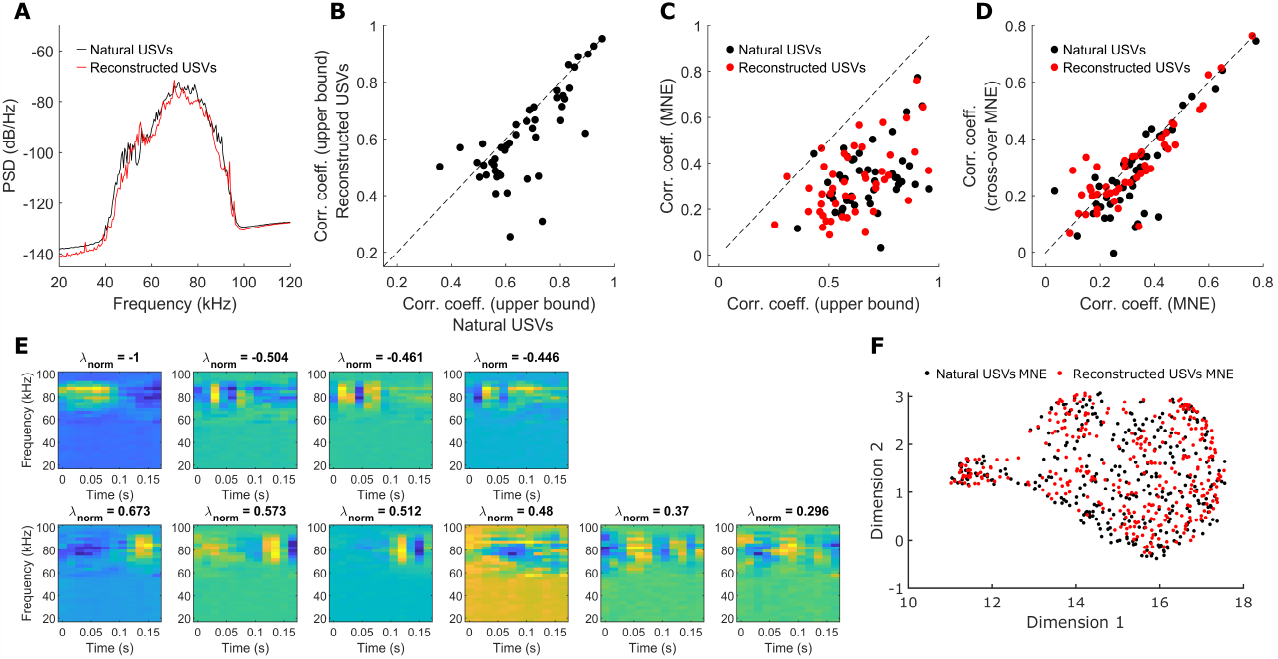
MNE model analysis characterising the response to BiWaveGAN-generated USVs. **(A)** Power spectra of natural and BiWaveGAN-reconstructed USVs. **(B)** Correlation coefficients between trials, indicating the upper bound on the correlation coefficients in response to the two stimulus types. Dashed line indicates a gradient of 1. **(C)** MNE model prediction correlation coefficients plotted as a function of the upper bound on the correlation coefficient achievable by the model. Dashed line indicates unity. **(D)** MNE model prediction correlation coefficients, where training and test data are swapped between natural and reconstructed USVs, plotted as a function of model performance using the within-class training and test data. **(E)** Example of one unit’s excitatory (top row) and inhibitory (bottom row) features learned by the MNE model using reconstructed stimuli. **(F)** UMAP projection onto two dimensions of the MNE model features learned with natural and reconstructed USVs.

The inherent variability of neural responses sets an upper limit on the correlation coefficient between responses to repeated presentations of a stimulus. This limit can be estimated by calculating the expected correlation coefficient indicating how consistently a stimulus evokes a neural response [***Hsu et al., 2004, Touryan et al., 2005***]. The expected correlation coefficients in response to both types of stimuli are approximately equal (Fig. 4**B**).

Maximum Noise Entropy (MNE) models were then trained for each unit, for natural USVs and reconstructed stimuli. An example of a unit’s excitatory and inhibitory features learned by the MNE model in response to reconstructed stimuli is shown in Figure 4**E**. The performance of each model, as measured by the correlation coefficient between the predicted response and the real response in the unseen test set, is shown in Figure 4**C**. These correlation coefficients are plotted against their upper bounds. One can see that the models’ prediction accuracies with the two classes of stimuli, natural and reconstructed USVs, are very similar.

The MNE models were then tested on the unseen test set of the other stimulus class, such that a model trained on natural USV stimulus activity was tested on a section of reconstructed USV stimulus activity, and vice versa. Model performance was compared to the performance using the test set of the same stimulus type (Fig.4**D**). Model performance is approximately equal between the two test data sets, further indicating that the natural and reconstructed USVs are equally suitable stimuli for estimating neurons’ receptive fields.

The similarity between MNE model features learned with the two stimuli was further shown (Fig. 4**F**) by projecting the features onto a low-dimensional manifold using the UMAP algorithm [***McInnes et al., 2020***].

Two measures of sparseness were then calculated to characterise the selectivity with which the neural population responds to the two types of stimuli [***Willmore and Tolhurst, 2001***]. The lifetime sparseness was calculated for each unit to indicate the degree of selectivity each unit has for different syllables within the two stimulus ensembles (Fig. 5**A**). Lifetime sparseness values near 100% indicate sparse coding, in which neurons respond strongly to only a few syllables, whereas values close to 0% indicate non-selective activity in response to all syllables in the stimulus ensemble. A Wilcoxon matched-pairs signed-rank test revealed a significant but small difference in the lifetime sparseness between the two stimuli (N=46 units; median and interquartile range (IQR) for natural USVs: 0.43 (0.32); for reconstructed: 0.40 (0.30); P=3.71e-6), with units showing a median decrease in sparseness of approximately 4% for the reconstructed stimulus.

**Figure 5.**
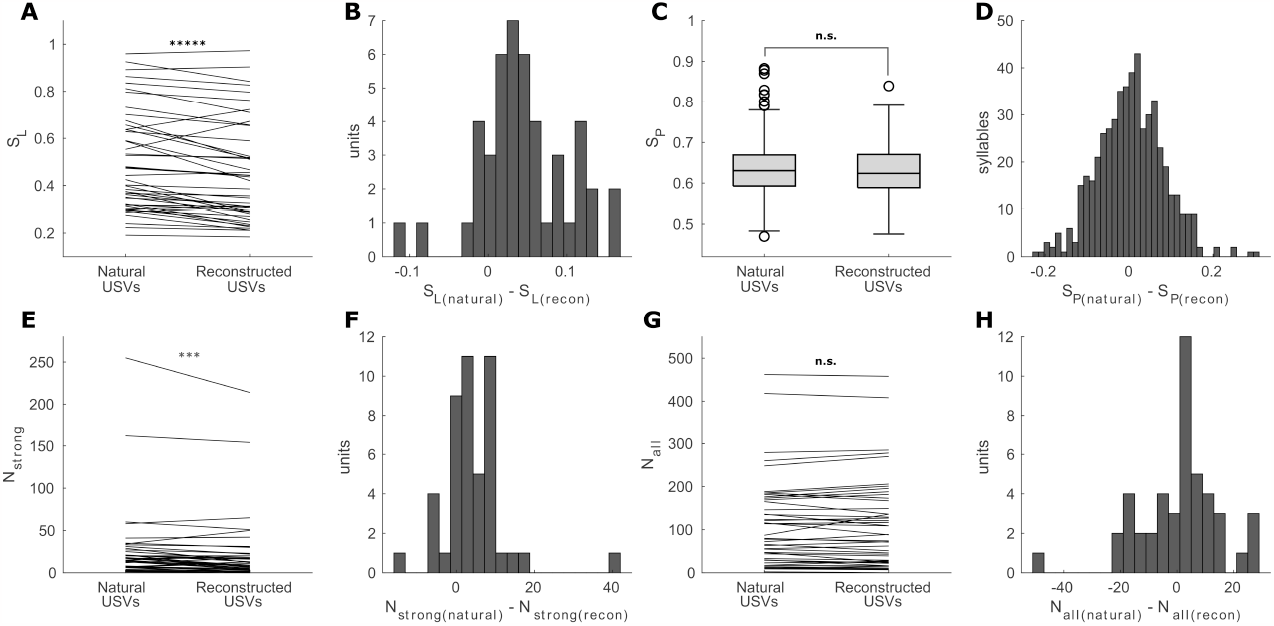
Sparseness of responses to natural and BiWaveGAN-generated USVs. **(A)** Lifetime sparseness calculated for each unit, for both natural USVs and BiWaveGAN-reconstructed stimuli. Data are presented as median and interquartile range (IQR). N=46 units; natural USV: 0.43 (0.32); reconstructed: 0.40 (0.30); Wilcoxon matched-pairs signed-rank test, P=3.71e-6. **(B)** A histogram of the difference in lifetime sparseness between the two sets of stimuli for each unit. **(C)** Population sparseness calculated for natural USVs and reconstructed stimuli. Data are presented as a box and whisker plot with median, first and third quartiles, and minimum and maximum values that are not outliers. Outliers are shown as circles (computed using the interquartile range). N=519 syllables; natural USVs: 0.63 (0.08); reconstructed: 0.62 (0.08); Wilcoxon matched-pairs signed-rank test, P=0.27. **(D)** A histogram shows the difference in population sparseness between the two stimuli. **(E)** Number of syllables evoking a high firing rate for each unit, defined as the response exceeding 50% of the maximum value of the PSTH to either stimulus type. N=46 units; natural USVs: 17 (20); reconstructed: 10 (16); Wilcoxon matched-pairs signed-rank test, P=1.50e-4. **(F)** A histogram shows the difference in the number of high firing rate syllables. **(G)** Number of syllables evoking any response for each unit, defined as the response exceeding 25% of the maximum value of the PSTH to either stimulus type. N=46 units; natural USVs: 79 (140); reconstructed: 81 (143); Wilcoxon matched-pairs signed-rank test, P=0.64. **(H)** A histogram shows the difference in the number of syllables evoking any response.

Population sparseness was then calculated to assess whether the two types of stimuli evoke more or less sparse codes: unlike lifetime sparseness, population sparseness characterises the responses of a set of neurons to each individual syllable rather than responses of each individual neuron to a set of syllables. Population sparseness values close to 0% indicate a dense code in which stimuli are encoded by a large proportion of the neural population, whereas population sparseness values close to 100% indicate a sparse code in which each stimulus is encoded by only a small fraction of the neural population. Note that as our units were recorded from multiple animals and only stimulus-responsive units are included in this analysis, this does not represent a true description of the full population response [***Hromádka et al., 2008***]. Rather, we use this metric only to compare the responses to the two types of stimuli. The population sparseness across all 519 syllables of both types of stimuli are shown in Fig. 5**C**. A Wilcoxon matched-pairs signed-rank test revealed no significant difference in population sparseness between them (median (IQR) for natural USVs: 0.63 (0.07); for reconstructed: 0.62 (0.08); P=0.27).

Lastly, to explain the apparent difference in findings between the two sparseness metrics, each unit’s response to each syllable was classified as either ‘strong’ or ‘weak’ by thresholding the PSTH. The maximum value of the PSTH (i.e., the maximum average firing rate) in response to either stimulus was taken as the maximal response. A response was classified as ‘strong’ if it exceeded 50% of this value, and as ‘weak’ if it exceeded 25% of this value. Figure 5**E** shows the number of syllables evoking ‘strong’ responses for each unit, and Figure 5**G** shows the sum of ‘strong’ and ‘weak’ responses. Although a significantly greater number of natural USV syllables evoke a strong response compared to reconstructed syllables (median (IQR) for natural USVs: 17 (20); for reconstructed: 10 (16); Wilcoxon matched-pairs signed-rank test, P=1.50e-4), both stimulus types evoke the same number of responses when including both strong and weak responses (median (IQR) for natural USVs: 79 (140); for reconstructed: 81 (143); Wilcoxon matched-pairs signed-rank test, P=0.64). This result indicates that the small difference in lifetime sparseness between the two stimuli is driven by a small number of very strong responses, which occur more often with the natural USV stimulus, resulting in increased lifetime sparseness. As population sparseness measures the distribution of responses over the population and is invariant to the magnitude of the response, this is shown to be the same between the two stimuli.

## Discussion

The use of natural stimuli to probe sensory systems has been shown to improve our ability to model and predict responses of single neurons [***Talebi and Baker, 2012, Laudanski et al., 2012***]. The improvement in model performance is associated with a distinct difference in the estimated receptive field structure [***Theunissen et al., 2000, David et al., 2004***]. At the population level as well there appears to be a marked difference in the responses to artificial and natural stimuli [***Hénaff et al., 2021***]. One can attribute this discrepancy to the inability of artificial stimuli to sample the full input space and thus biasing computational models [***Sharpee, 2013***]. In the case of auditory stimuli, artificial sounds such as pure tones are unable to capture the complex correlation structure that is present in natural vocalisations or recordings of the natural environment. Whilst efforts have been made to devise artificial stimuli that can overcome this limitation [***Depireux et al., 2001, Atencio and Schreiner, 2012***], they remain inferior to using natural vocalisations [***Laudanski et al., 2012***].

We aim to bridge the gap between artificial and natural stimuli by providing a method to systematically generate naturalistic vocalisations with near-identical properties to vocalisations produced naturally. We show that these stimuli evoke comparable neural responses, that the receptive fields obtained using synthesised stimuli have the same predictive power as those obtained with natural stimuli, and that these receptive-field features form a single cluster when projected in the lowdimensional UMAP space.

By exposing the latent space used to generate these vocalisations, it is also possible to sample the sensory input space in a more principled manner by drawing homogeneously spaced samples from the latent space. This could improve the performance of existing models of sensory processing by reducing the bias introduced by *ad hoc* stimulus selection. Additionally, it is possible to smoothly interpolate between different vocalisations (Fig. 2) which can be used to study decision making in a more natural context [***Sainburg et al., 2022***]. Similar methods exist for the generation of vocalisations [***Sainburg et al., 2020, Arneodo et al., 2021***], however, these models generate spectrograms whose phase must be reconstructed to produce waveforms, while our model produces waveforms directly.

Generative models like BiWaveGAN can be used to discern the structure of animal communications in an unsupervised, unbiased manner. While valuable in their own right, these models could be particularly useful for researchers relying on the playback of vocalisations as stimuli to study the neurological and behavioural responses of animals. Using such a model in this manner eliminates the need to record large volumes of vocalisations from animals, which can be time consuming in the laboratory and difficult in the wild [***Chabout et al., 2017***]. Once trained, the model can efficiently generate large volumes of diverse samples which can be continuously varied via interpolation in the latent space, and clusters of highly similar but not identical samples can be produced by sampling nearby points in latent space. In addition, such models are not restricted to any one type of animal but could include multiple species. This would allow for interpolation between vocalisations of different species to study the importance of ethological relevance in auditory encoding.

## Methods and Materials

### Dataset

Samples of ultrasonic vocalisations were acquired from data recorded by ***Van Segbroeck et al***. [***2017***]. Briefly, the data consist of spontaneous vocalisations from two inbred strains (C57Bl/6J and DBA/2J) recorded at a sample rate of 250 kHz. These recordings were then processed and segmented by segmentation software (MUPET [***Van Segbroeck et al., 2017***], using default parameters). This produced a total of 31475 individual syllables which were randomly divided into a training set of 28329 syllables and a held-out test set of 3164 syllables (90% and 10% of the total data, respectively). Figure S1 shows a sample of the data visualised as spectrograms. Before training the model, the length of individual audio samples was adjusted to accommodate the WaveGAN architecture which requires equal sample length. The length was chosen to be 2^15^ = 32768 samples, which equates to approximately 130 ms when sampled at 250 kHz. Samples were either zero-padded or truncated to achieve the desired length.

### Model Architecture

Given a dataset of samples *X* = {*x*_1_, *x*_2_, …, *x*_*N*_} where *x*_*i*_ is the waveform of an individual USV syllable, we assume these data have been drawn from an unknown probability distribution *p*_*X*_(*x*). The task of the generative model is to learn a distribution 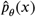 that approximates the true underlying distribution *p*_*X*_(*x*) of the data. Here *θ* represents the parameterising variables of the model, and training the generative model means iteratively optimising *θ* such that 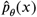 approximates *p*_*X*_(*x*). Generating new data then simply requires sampling data points 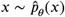 from the model distribution.

We construct the generative model following the WaveGAN architecture [***Donahue et al., 2019***]. The generator consists mainly of 1D transpose convolutional layers. In these layers the input is upsampled by a factor of 4 with new elements set to zero; this is followed by a convolution with a kernel of length 25 whose weights are to be optimised during training. Each layer is followed by a *ReLU* (Rectified Linear Unit) activation function, except the last layer which is followed by a *tanh* activation function (Table 1, left). The output of these layers is twice as long as the input, therefore by concatenating several layers the generator can expand the latent vector into a full waveform.

**Table 1.** Architecture tables for BiWaveGAN generator (left) and encoder (right). All Conv1D and TransConv1D layers have a kernel of length 25. All LeakyReLU activations have a negative slope of 0.2. *m* is the batch size, *l* is the dimension of latent space and *d* is a parameter controlling model size.

| $G$ | Operation | Output Shape | Activation | $E$ | Operation | Output Shape | Activation |
| --- | --- | --- | --- | --- | --- | --- | --- |
| 0 | Input $z$ batch | $(m, l)$ | | 0 | Input $x$ batch | $(m, 1, 32768)$ | |
| 1 | Reshape | $(m, 1, l)$ | | 1 | Conv1D(stride=2) | $(m, 2d, 16384)$ | LeakyReLU |
| 2 | Fully Connected | $(m, 1, 512d)$ | ReLU | 2 | Conv1D(stride=4) | $(m, 4d, 4096)$ | LeakyReLU |
| 3 | Reshape | $(m, 32d, 16)$ | | 3 | Conv1D(stride=4) | $(m, 8d, 1024)$ | LeakyReLU |
| 4 | TransConv1D(stride = 4) | $(m, 32d, 64)$ | ReLU | 4 | Conv1D(stride=4) | $(m, 16d, 256)$ | LeakyReLU |
| 5 | TransConv1D(stride = 4) | $(m, 16d, 256)$ | ReLU | 5 | Conv1D(stride=4) | $(m, 32d, 64)$ | LeakyReLU |
| 6 | TransConv1D(stride = 4) | $(m, 8d, 1024)$ | ReLU | 6 | Conv1D(stride=4) | $(m, 32d, 16)$ | LeakyReLU |
| 7 | TransConv1D(stride = 4) | $(m, 4d, 4096)$ | ReLU | 7 | Reshape | $(m, 1, 512d)$ | |
| 8 | TransConv1D(stride = 4) | $(m, 2d, 16384)$ | ReLU | 8 | Fully Connected | $(m, 1, l)$ | |
| 9 | TransConv1D(stride = 2) | $(m, 1, 32768)$ | Tanh | 9 | Reshape | $(m, l)$ | |

The discriminator (or ‘critic’) consists of three sub-networks. First, a waveform *x* is processed by ***D***_*x*_, and a latent vector *z* is processed by ***D***_*z*_. ***D***_*x*_ consists of 1D convolutional layers with kernels of length 25 and a stride of 4 (except for the first convolutional layer, which has a stride of 2). Each layer is followed by a *LeakyReLU* activation function with a negative slope of 0.2 and then a phase shuffle operation (Table 2). A phase shuffle operation randomly perturbs the phase of its input by a random amount. This is necessary because the up-sampling performed by the generator causes artefacts in the generated audio that occur with a particular phase which the discriminator can exploit. By permuting the phase, the discriminator is made to be roughly invariant to the phase of the input waveform [***Donahue et al., 2019***]. ***D***_*z*_ consists of three convolutional layers with a kernel length of 1 and a stride of 1, followed by a *LeakR****E****LU* activation function.

**Table 2.** Architectures of ***D***_*x*_, ***D***_*z*_ and ***D***_*joint*_ in a BiWaveGAN discriminator, where ***D***_*z*_ has a depth of 3 and ***D***_joint_ has a depth of 3. Stride = 1 for all convolutions unless otherwise stated. *m* is the batch size, *l* is the dimension of the latent space, *d* is a parameter controlling model size, *n* controls the amount of PhaseShuffle that is applied, and *f* controls the number of filters used in the convolutions of ***D***_*z*_ and ***D***_joint_. In our model, the values *d* = 32, *n* = 2 and *f* = 512 were used.

| $D_x$ | Operation | Output Shape | Activation |
| --- | --- | --- | --- |
| 0 | Input $x$ | $(m, 1, 32768)$ | |
| 1 | Conv1D(kernel length = 25, stride = 2) | $(m, d, 8192)$ | LeakyReLU |
| 2 | PhaseShuffle($n$) | $(m, d, 8192)$ | |
| 3 | Conv1D(kernel length = 25, stride = 4) | $(m, 2d, 2048)$ | LeakyReLU |
| 4 | PhaseShuffle($n$) | $(m, 2d, 2048)$ | |
| 5 | Conv1D(kernel length = 25, stride = 4) | $(m, 4d, 512)$ | LeakyReLU |
| 6 | PhaseShuffle($n$) | $(m, 4d, 512)$ | |
| 7 | Conv1D(kernel length = 25, stride = 4) | $(m, 8d, 128)$ | LeakyReLU |
| 8 | PhaseShuffle($n$) | $(m, 8d, 128)$ | |
| 9 | Conv1D(kernel length = 25, stride = 4) | $(m, 16d, 32)$ | LeakyReLU |
| 10 | Conv1D(kernel length = 25, stride = 4) | $(m, 32d, 15)$ | LeakyReLU |
| 11 | Conv1D(kernel length = 25, stride = 4) | $(m, 32d, 1)$ | LeakyReLU |

| $D_z$ | Operation | Output Shape | Activation |
| --- | --- | --- | --- |
| 0 | Input $z$ batch | $(m, l)$ | |
| 1 | Reshape | $(m, l, 1)$ | |
| 2 | Conv1D(kernel length = 1) | $(m, f, 1)$ | LeakyReLU |
| 3 | Conv1D(kernel length = 1) | $(m, f, 1)$ | LeakyReLU |
| 4 | Conv1D(kernel length = 1) | $(m, f, 1)$ | LeakyReLU |

**Table 2.** Architectures of $D_x$ , $D_z$ and $D_{joint}$ in a BiWaveGAN discriminator, where $D_z$ has a depth of 3 and $D_{joint}$ has a depth of 3.
| $D_{joint}$ | Operation | Output Size | Activation |
| --- | --- | --- | --- |
| 0 | Concat($D_x(x), D_z(z)$) | $(m, 32d + f, 1)$ | |
| 1 | Conv1D(kernel length = 1) | $(m, f, 1)$ | LeakyReLU |
| 2 | Conv1D(kernel length = 1) | $(m, f, 1)$ | LeakyReLU |
| 3 | Conv1D(kernel length = 1) | $(m, 1, 1)$ | |
| 4 | Reshape | $(m, 1)$ | |

Afterwards, the two feature maps produced by ***D***_*x*_ and ***D***_*z*_ are concatenated and fed into the final network ***D***_joint_ which produces the final output. ***D***_joint_ also consists of three convolutional layers with a kernel length of 1 and a stride of 1, followed by a *LeakR****E****LU* activation function, except for the final layer which has no activation function. Therefore ***D***(*x*, z) = ***D***_*joint*_(Concat(***D***_*x*_(*x*), ***D***_*z*_(*z*)).

Similarly, the encoder network used to sample the model distribution, 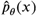, consists of onedimensional convolutions followed by *LeakyReLU* activation functions. The kernel lengths were chosen to be the inverse of the generator’s. The stride of each convolution layer is 4, except for the first convolution layer where it is 2. Unlike the discriminator network, there are no phase shuffle operations (Table 1, right).

Table 1 shows the architectures of ***G*** and ***E***, placed side by side for comparison. Notice how ***E*** has a roughly inverse architecture to ***G***, each TransConv1D in ***G*** having a corresponding Conv1D in ***E***, since we want ***G*** and ***E*** to be approximate inverses. Figure 6 displays the full architecture of all three models.

**Figure 6.**
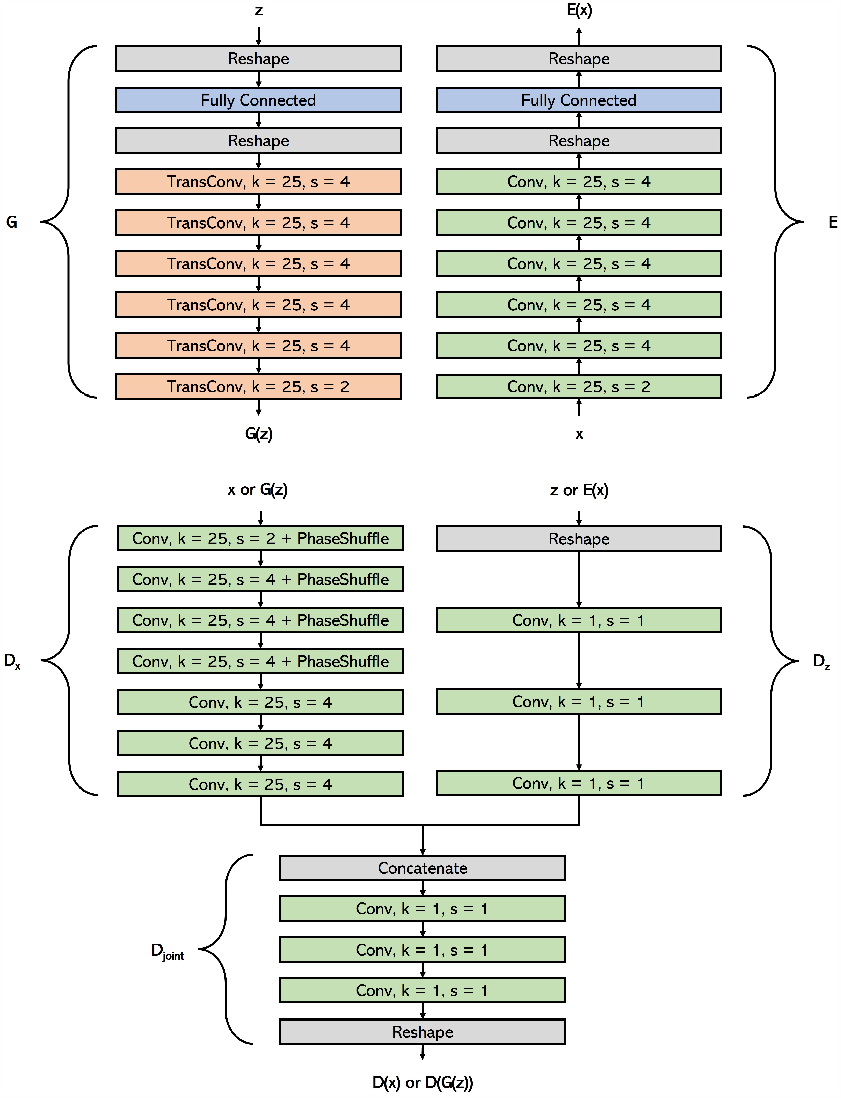
BiWaveGAN model architecture. BiWaveGAN consists of three networks: the generator (***G***), the encoder (***E***), and the discriminator or critic (***D***) which consists of the sub-networks ***D***_*x*_, ***D***_*z*_ and ***D***_joint_. Conv and TransConv layers represent 1D Convolutional and Transpose Convolutional layers, where *k* is the length of the kernel, and *s* is the stride of the convolution or transpose convolution. During training ***D*** learns to approximate the Wasserstein-1 distance between the distributions of real data-vector pairs, (*x*, ***E***(*x*)), and synthetic pairs, (***G***(*z*), z) while ***G*** and ***E*** learn to produce data-vector pairs which minimise this distance. After training, ***G*** can be fed latent vectors to produce synthetic waveforms, and ***E*** can be fed waveforms to infer their latent representations.

The dataset, model training script and model weights can be accessed via GitLab ^1^.

### Model Training

As stated previously, GANs consist of a generator network, ***G***, and a discriminator network, ***D***, that work against each other during the training process. ***G*** maps latent vectors z to artificial data points ***G***(*z*), while ***D*** takes inputs from the data domain and outputs a value in [0, 1]. ***D***(*x*) can be interpreted as the probability according to ***D*** that *x* comes from the real data distribution, so a perfect discriminator satisfies ***D***(*x*) = 1 for all real data *x* and ***D***(***G***(*z*)) = 0 for all fake data ***G***(*z*). Let *p*_*X*_ be the true underlying distribution of our data and *p*_*z*_ be the distribution of our latent variable (defined as either a Gaussian or uniform distribution). In the original GAN formulation, the training objective is as follows [***Goodfellow et al., 2014***]:

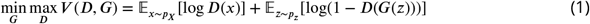

This means that ***D*** attempts to maximise its output for real data and minimise its output for fake data generated by ***G***, while ***G*** attempts to maximise ***D***’s output for the fake data that it generates. The weights of ***G*** and ***D*** can be optimised via iterative methods, e.g., stochastic gradient descent.

Due to the adversarial nature of their training, GANs can be unstable and suffer from a phenomenon known as mode collapse in which the generator only learns to generate a subset of the data distribution, resulting in less variety in the generated data than in the real data. To alleviate these shortcomings, ***Arjovsky et al***. [***2017***] developed a variant known as Wasserstein GAN or WGAN.

It can be shown that the original GAN optimisation problem in Eq. (1) is equivalent to minimising the Jensen-Shannon divergence between the distributions *p*_*X*_ and *p*_***G***(*z*)_ (i.e., the real and fake data). However, the authors argue that it is more beneficial to optimise the Wasserstein-1 distance instead, which can be expressed as follows:

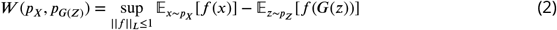

Note that *f* maps data points (real or fake) to any real number and ‖*f*‖ _*L*_ ≤ 1 means that *f* is 1-Lipschitz continuous, i.e., its derivative (assuming *f* is differentiable) is bounded by 1: ∇_*x*_*f* (*x*) ≤ 1 for all *x*. Thus the Wasserstein-1 distance is the *supremum* of 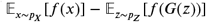 over the set of 1-Lipschitz continuous functions.

Therefore in a WGAN we replace Eq. (1) with the following optimisation objective:

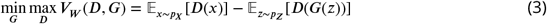

Note that here ***D*** is playing the role of *f* in Eq. (2), and its job is no longer to distinguish between real and fake inputs but to assist in computing the Wasserstein-1 distance by maximising *V*_*W*_ (***D, G***)—hence it is often referred to as a ‘critic’ rather than a ‘discriminator’. Therefore ***D***’s outputs are not restricted to the range [0, 1] but can be any real number.

Recall that the definition of the Wasserstein-1 distance requires *f* to be 1-Lipschitz continuous, but ***D*** is in practice a neural network which does not necessarily satisfy this property. To enforce the 1-Lipschitz continuity of ***D, Gulrajani et al***. [***2017***] propose Wasserstein GAN with Gradient Penalty (WGAN-GP). This means adding a term onto the loss function of ***D*** that punishes ***D*** when its gradients are larger than one.

The encoder, ***E***, is trained adversarially alongside ***G*** and ***D***. Furthermore, we must modify our discriminator so that it accepts as input both a data point and a latent vector. During training, instead of showing the discriminator *x* or ***G***(*z*), we show it the tuples (*x*, ***E***(*x*)) or (***G***(*z*), z). The adversarial objective of a BiGAN is as follows:

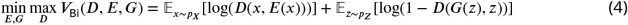

Note the similarities between this and Eq. (1) for regular GANs, and that ***E*** and ***G*** have the same objective. It can be shown that in order for ***G*** and ***E*** to ‘fool’ a perfect discriminator at a point in the joint space (*x*, z) they must satisfy *x* = ***G***(***E***(*x*)) and z = ***E***(***G***(*z*)), i.e., ***E*** and ***G*** must learn to be inverse to each other, as we desire.

Our aim is to adapt WaveGAN into a BiGAN. Since WaveGAN is a WGAN-GP model, we must first synthesise the optimisation objectives and training procedures of WGAN-GP and BiGAN. By doing this we get the stable training and improved convergence properties of WGAN-GP as well as the latent-representation enabled by BiGAN.

Looking at Equations (3) and (4) we see that they can be combined into:

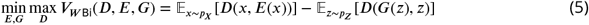

Training a model around this optimisation objective can then be done by accommodating the encoder into the WGAN-GP training procedure as can be seen in Alg. 1.

The model was trained using an Adam optimiser [***Kingma and Ba, 2017***] with a learning rate of 0.0001 for 150,000 iterations with a batch size of 64. Inputs to the generator are sampled from a Uniform[-1,1] distribution. The model was trained on an NVIDIA Titan Xp GPU.

### Surgical preparation

All procedures were carried out under the terms and conditions of licences issued by the UK Home Office under the Animals (Scientific Procedures) Act 1986. Extracellular recordings were made in the auditory cortex of adult female C57BL/6J mice (N=6, aged 6 to 11 weeks). Animals were anaesthetised using a mixture of fentanyl, midazolam and medetomidine (0.05, 5 and 0.5 mg/kg, respectively). A midline incision was made over the dorsal surface of the cranium and the right temporalis muscle was partially resected. Stereotaxic coordinates were used to locate the right auditory cortex, using a rostro-caudal coordinate of 70% bregma-lambda and a dorso-ventral coordinate of bregma -2.2 mm (the lateral coordinate being determined by the surface of the skull). A steel headplate comprising a bent piece of flat bar (approximately 5 mm x 30 mm) was attached to the dorsal surface of the skull using a combination of tissue adhesive (Histoacryl) and dental cement (Kemdent works, Swindon, UK). This was subsequently used in combination with a magnetic stand to secure the animal in place for the remainder of the surgery and recordings.

#### Algorithm 1 Training a Bidirectional WGAN-GP via minibatch gradient descent. Parameters are

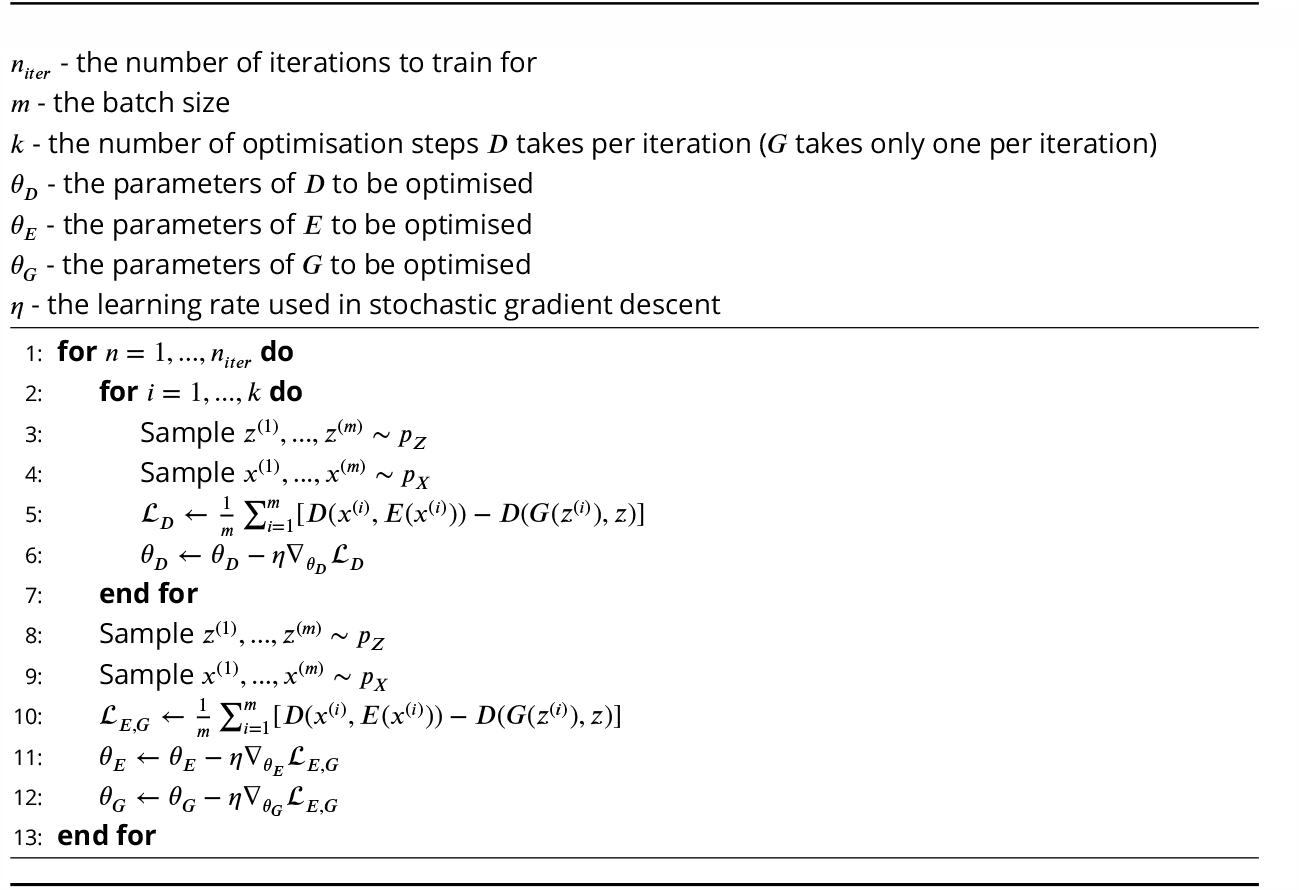

A small (*ϕ*=2mm) craniotomy was made over the auditory cortex using a dental drill and burr, and a small hole was made in the dura. The surface of the brain was protected from desiccation by regular application of silicone oil. A machine screw (M1 x 2 mm) was inserted into the skull approximately over the contralateral motor cortex to act as a ground for the probe. The animal was then transferred to an acoustic chamber, and once again secured by fixation of the headpost in a magnetic stand. Body temperature was maintained throughout the duration of the experiment using activated Deltaphase isothermal pads (Braintree Scientific, Braintree, USA).

### Syllable preprocessing and Stimulus Presentation

Auditory stimuli consisting of 519 syllables of natural and BiWaveGAN-reconstructed USVs were generated. Syllables were first filtered by 6th-order Chebyshev Type II filters (40 kHz highpass and 100 kHz lowpass), and noise was reduced using the Noise reduction tool in Audacity ^2^. Syllables were then concatenated, separated by 100ms of silence. Natural USV and reconstructed USV stimuli were played alternately for a total of at least 20 repetitions of each stimulus type. Stimuli were presented using an Avisoft UltraSoundGate Player 116 in combination with an Avisoft Vifa ultrasonic speaker (Avisoft Bioacoustics, Glienicke, Germany). Signal intensity was set such that the peak intensity did not exceed 85 dB SPL. For clarity, all spectrograms are displayed thresholded at 2 standard deviations above the mean power.

### Electrophysiological recording

The experimental setup consisted of a silicon multi-electrode probe (single-shank, 32-channel polytrode, Neuronexus) connected to a Neuronexus Smartbox Pro data acquisition system which amplified the signals and digitised them at a sampling rate of 30 kHz. The conductive area of the recording sites measured 177*μm*^2^. Spike sorting was done offline using an automated algorithm (Kilosort3 [***Pachitariu et al., 2023***]). Units detected by Kilosort3 were subsequently manually inspected and curated using Phy [***Rossant and Harris, 2013***]. Units were considered to be well-isolated single units if their spike waveforms displayed consistent shapes and their autocorrelograms exhibited clear refractory periods. Units were then classified as being auditory by manual inspection of raster plots to identify stimulus-locked spikes. Units that were lost (e.g., due to electrode drift) resulting in recordings of fewer than 10 trials of stimulus presentation were excluded.

### Receptive field characterisation

The receptive fields of single units were characterised using the Maximum Noise Entropy (MNE) method [***Fitzgerald et al., 2011***]. Briefly, this method comprises fitting a model that describes the probability of a spike as a function of a stimulus, **s**, as *P* (*spike*|**s**) = (1 + *exp*(*a* + **s***h* + **s**^*T*^ ***J* s**)^−1^, where *a, h* & ***J*** are parameters optimised via gradient descent. These parameters are determined such that the predicted firing rate, spike-triggered average and spike-triggered covariance match those in the observed data. The specific parameters used for the spectro-temporal representation of the stimulus data were chosen through bioacoustical analysis, as detailed in our previous work [***Lu et al., 2023***].

The MNE model for each unit was trained with 80% of the stimulus-response data. The remaining 20% of the data were used to test the model. The training data were randomly extracted from the full recording without replacement to minimise the effects of any non-stationarity of the data. After training the MNE model for each unit, the model was used to predict a response to the test stimulus. The correlation coefficient between the model predictions and the recorded neural activity was calculated to assess the performance of the model.

Neural activity of a single unit is not perfectly correlated between repeated presentations of the stimulus; this affects how the performance of a model based on its correlation coefficient with recorded data is interpreted. To account for this inherent variability in the neural activity, we estimate the expected correlation coefficient between the responses to repeated presentations of the same stimulus in the test set [***Hsu et al., 2004, Touryan et al., 2005***]. The expected correlation coefficient 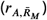 is defined as the correlation between the true or actual firing rate, *A*, and the firing rate 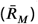 as measured by averaging over *M* repeats. As detailed in [***Hsu et al., 2004***], we can calculate 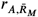 using the following equation:

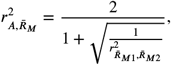

where 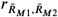 is the correlation between 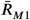 and 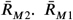 is the firing rate estimated by averaging over *M/*2 repetitions and 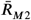 is the firing rate estimated by averaging over the other *M/*2 repetitions of the neural response.

The correlation coefficients between the model prediction and the test set were then scaled by the expected correlation of the test set.

### Feature comparison

To compare the similarity between MNE features estimated from neurons’ responses to real or reconstructed USVs, we projected the modulation power spectra of the features into a 2D latent space using the UMAP algorithm [***McInnes et al., 2020***] with parameters *n*_*neighbors* = 256, *min*_*dist* = 0, *n*_*dim* = 2 and *metric* = ′*euclidean*′. Modulation power spectra were obtained by computing the two-dimensional Fourier transform of each spectrogram as described in ***Atencio and Sharpee*** [***2017***].

### Sparseness Measurement

Lifetime and population sparseness were calculated using the definition for sparseness given in [***Vinje and Gallant, 2000***]:

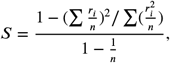

In the case of lifetime sparseness, *r*_*i*_ represents the average firing rate of a single unit in response to the *i*^*th*^ stimulus in the ensemble of n individual stimuli; in the case of population sparseness, *r*_*i*_ represents the average firing rate of the *i*^*th*^ unit in the population of n units, during the presentation of a single stimulus. Lifetime sparseness thereby characterises the response of an individual unit to a set of stimuli, whereas population sparseness characterises the response of a whole population of units to a single stimulus. Lifetime sparseness values near 0 indicate a dense code, in which the neuron responds with equal firing rate to all stimuli in the ensemble, whereas values close to 1 indicate a sparse code in which the neuron responds highly selectively to only a few stimuli. Similarly, population sparseness values close to 0 indicate a dense code in which a large proportion of the population responds to a stimulus, whereas values close to 1 indicate that only a small fraction of the population responds to the stimulus.

## Funding

This work was funded by Biotechnology and Biological Sciences Research Council grant “How do auditory cortical neurons represent ethologically relevant natural stimuli? Characterizing stimulus feature selectivity and invariance” (BB/N008731/1).

## Supplementary

**Figure S1.**
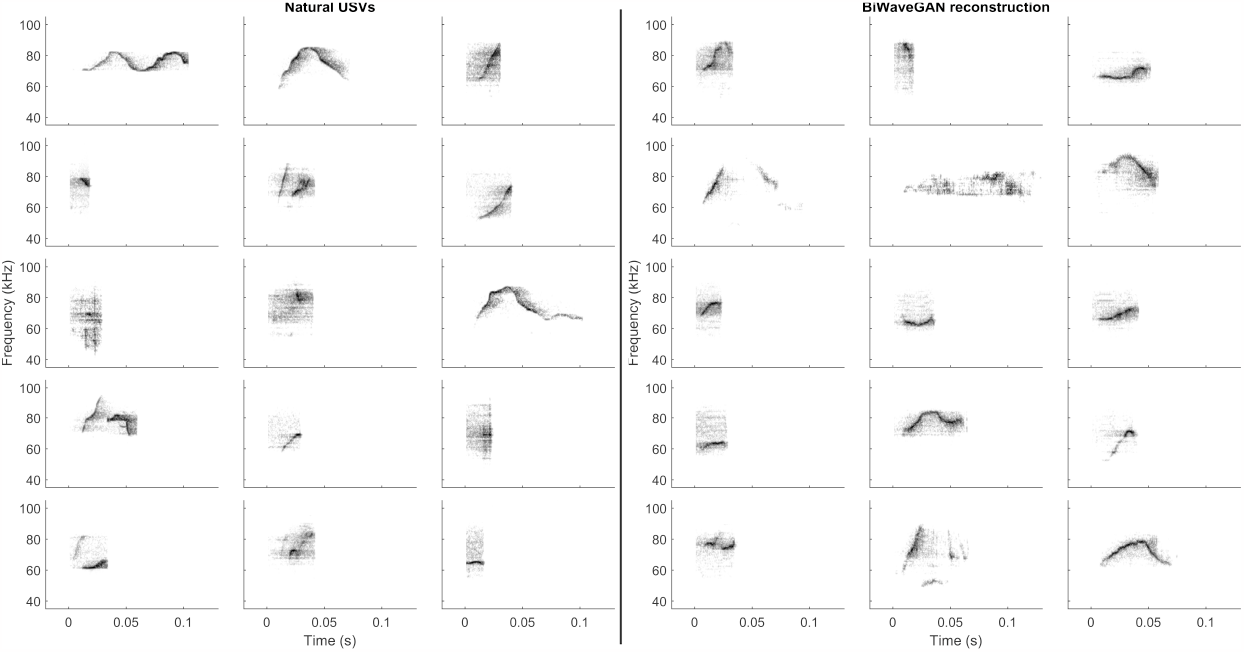
Spectrograms of a random sample of natural USVs and USVs generated by the BiWaveGAN model. The model is capable of generating a wide variety of USV types encountered in the training data.

## Footnotes

1 https://gitlab.com/kozlovlabcode/biwavegan

2 https://audacity.com

## Notes

https://gitlab.com/kozlovlabcode/biwavegan

### Competing Interest Statement

The authors have declared no competing interest.

